# A Divide-and-Conquer Approach to Large-Scale Evolutionary Analysis of Single-Cell DNA Data

**DOI:** 10.1101/2024.04.28.591536

**Authors:** Yushu Liu, Luay Nakhleh

## Abstract

Single-cell sequencing technology is producing large datasets, often containing thousands or even tens of thousands of single-cell genomic data points from an individual patient. Evolutionary analyses of these data sets help uncover and order genetic variants in the data as well as elucidate mutation trees and intra-tumor heterogeneity (ITH) in the case of cancer data sets. To enable such large-scale analyses computationally, we propose a divide-and-conquer approach that could be used to scale up computationally intensive inference methods. The approach consists of four steps: 1) partitioning the dataset into subsets, 2) constructing a rooted tree for each subset, 3) computing a representative genotype for each subset by utilizing its inferred tree, and 4) assembling the individual trees using a tree built on the representative genotypes. Besides its flexibility and enabling scalability, this approach also lends itself naturally to ITH analysis, as the clones would be the individual subsets, and the “assembly tree” could be the mutation tree that defines the clones. To demonstrate the effectiveness of our proposed approach, we conducted experiments employing a range of methods at each stage. In particular, as clustering and dimensionality reduction methods are commonly used to tame the complexity of large datasets in this area, we analyzed the performance of a variety of such methods within our approach.

## 1 Introduction

Copy number alterations (CNAs) refer to changes in the number of copies of DNA segments, including amplifications and deletions of genomic sections [5]. These alterations are crucial in identifying various types of cancers characterized by uncontrolled cell growth and invasion of other body parts [18,1,3]. At a genetic level, CNAs in critical genes can cause either over-expression or loss of gene functions, ultimately leading to the initiation and progression of cancers [2,21]. Therefore, studying CNAs can provide insights into understanding the underlying mechanisms of tumorigenesis, which may help identify therapy targets in cancer treatment [8].

The recent advancement of single-cell sequencing technology has revolutionized our understanding of how cancer evolves [35,11]. It enables the sequencing of hundreds to thousands of cells from a single tumor, e.g., [19,9], offering an unparalleled level of detail. However, single-cell DNA sequencing data (scDNAseq) also presents significant computational challenges [14].

Divide-and-conquer approaches are common in phylogenetics and have enabled the inference of phylogenies on very large datasets [36]. These approaches typically divide the large set of taxa into overlapping subsets, infer trees on the subsets, and then merge them using “supertree” methods (methods that glue small trees into a large one). Such an approach is not directly applicable to large scDNAseq datasets since such datasets have very little signal in them to enable accurate inference of the subtrees. In this paper, we present a different divide-and-conquer approach for the evolutionary analysis of large scDNAseq data. The approach has four steps. First, the dataset is partitioned into subsets of individual cells. Second, we infer a tree on each subset. Third, we compute a representative genotype for each subset using its tree. Finally, we infer an “assembly tree” on the representative genotypes to glue the subtrees. It is important to note here that while tree inferred in the second step might not be accurate (due to low signal, as we mentioned above), we show that ancestral genotype inference is robust to tree error, and that obtaining such representative genotypes yields better results than computing consensus genotypes. This approach is flexible and allows for incorporating a diverse array of methods for each step, from phylogenetically informed partitioning to various tree construction and genotype inference techniques. Additionally, the approach could be applied recursively on the subsets, thus controlling for the sizes of the individual subsets to match the computational requirements of the employed tree inference methods.

As clustering with or without dimensionality reduction is commonly used in large-scale scDNAseq datasets [9,31], we evaluated the performance of three clustering methods, Louvain, K-means, and hierarchical clustering, as these were the methods used in studies such as [23,15,37,17] and two dimensionality reduction techniques, PCA (Principal Component Analysis) and t-SNE (t-distributed Stochastic Neighbor Embedding), for the first step of our approach. Dimensionality reduction has been used in several studies, e.g., [6,22,4,32].

For the third step, we evaluated the use of consensus genotypes and compared that to our proposed ancestral genotype inference. We found the latter to provide more accurate results. Finally, for inferring an “assembly tree” that glues the subtrees together, we used our recently developed method, NestedBD [20], to infer a tree on the copy number profiles (the representative genotypes).

We evaluated the approach on both simulated and biological data. We show that the choice of a clustering method could make a big difference in the obtained results. Nevertheless, by choosing the right partitioning method, the approach yields very good results and scales up analyses to very large datasets.

## 2 Methods

### 2.1 A Divide-and-conquer Approach

An outline of the approach is shown in Fig. 1. The approach consists of four steps: 1) Partitioning the dataset into subsets through clustering; 2) Constructing a rooted tree for each subset; 3) Inferring ancestral genotypes for the roots of these trees to serve as the cluster representative profiles (CRPs); and 4) Assembling a unified tree from these subsets based on the inferred ancestral genotypes.

**Fig. 1.**
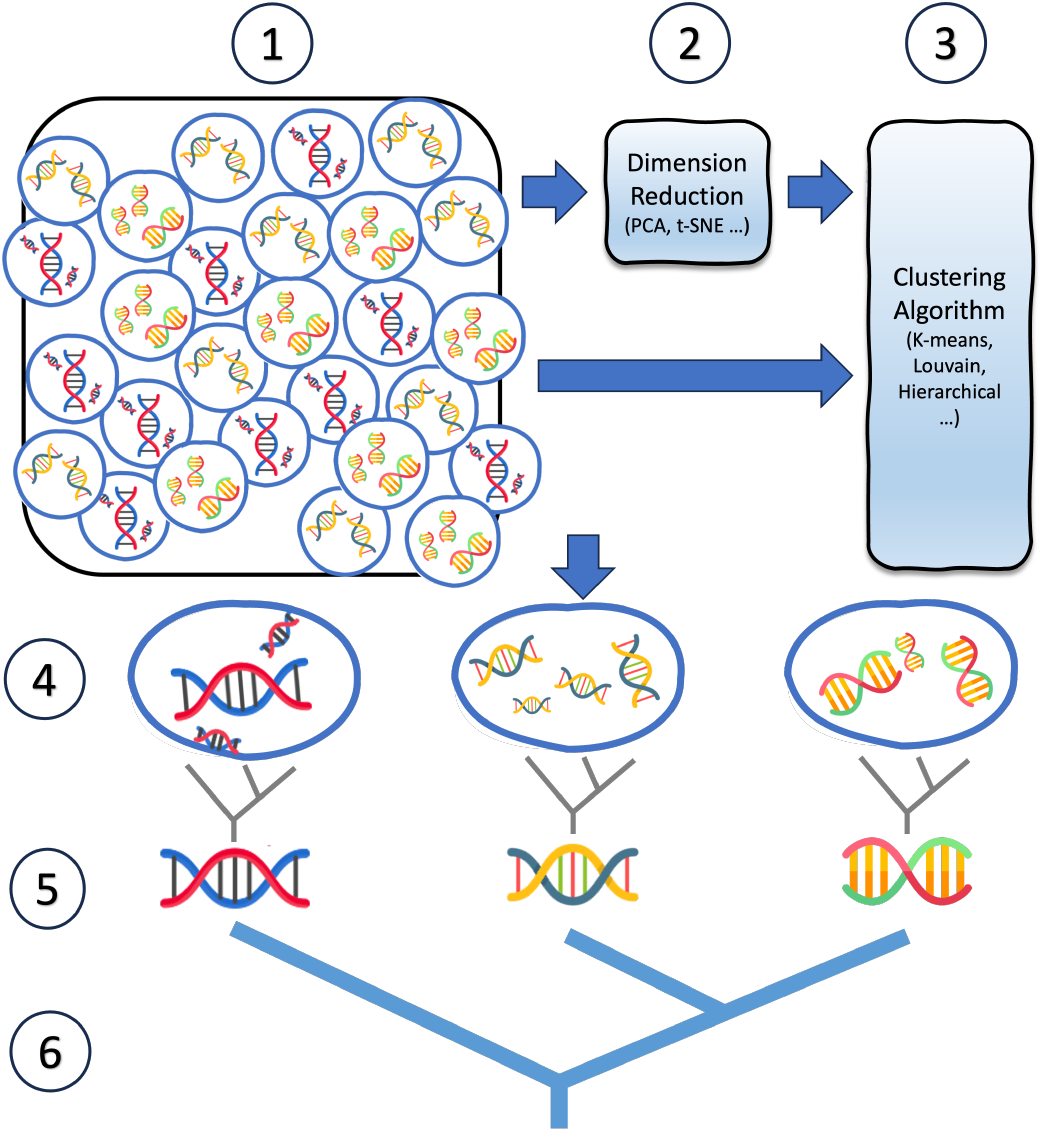
Divide-and-conquer evolutionary analysis approach. (1) Single-cell DNA sequencing allows for the generation of the genomes of thousands of cells. Copy number profiles (CNPs) would then be inferred for these genomes, resulting in one CNP per cell. (2) Dimensionality reduction could be applied to the data. (3) The data, in its original dimensions or after dimensionality reduction, is partitioned using, for example, a clustering method of choice. (4) A number of subsets, or clusters, each with a much smaller number of CNPs than the original data set, are obtained. (5) A cluster-representative profile, or CRP, of each cluster is computed by inferring ancestral genotypes at the roots of tree built from the CNPs in the corresponding cluster. (6) A tree on the CRPs is obtained and can be viewed as a mutation tree on the full data set, where the individual cells in a cluster are attached to the leaf labeled by the corresponding cluster’s CRP, or a rooted tree computed on the CNPs within the cluster is attached to that leaf.

We experimented with a variety of methods for each step, from clustering to tree construction. It is important to emphasize that our objective is not to declare a definitive best method for each step but rather to illustrate the approach’s flexibility and broad applicability across diverse datasets and analytical challenges. To this end, we applied two dimensionality reduction techniques, PCA and t-SNE, to the scDNA copy number data. Following dimensionality reduction, we clustered the cells using three algorithms, Louvain, K-means, and hierarchical clustering.

We introduce an ancestral profile (AP) based approach for generating cluster-representative profiles (CRPs) (step (5) of Fig. 1). This approach calculates the ancestral profile at the root of the evolutionary tree constructed from all profiles in a cluster, thereby representing the common ancestor of all profiles within that cluster. Finally, we inferred trees on the CRPs (step (6) of Fig. 1) using NestedBD [20].

### 2.2 Simulation Protocol

As the first step, we simulated an ultra-metric tree under a birth-death model with equal birth rate and death rate with 1k, 5k, and 10k extent leaves using the simulator described in [33]. We set the root node as a diploid without any CNA, and each leaf node represents a single cell sample from the patient. The internal nodes represent the cells that existed in the past and were not sampled. The branch lengths of the simulated trees are scaled to ensure all simulated trees have a height of 1, keeping the expected number of events uniform across simulation settings, irrespective of the number of sampled cells. Then, the simulator described in [20] is used to simulate copy number profiles given the tree topology, with simulation parameters chosen based on their ability to simulate CNAs that best resembled those observed in the biological data sets studied as in [20].

Then, to simulate CNA events along the tree, we first determine the number of CNAs of the branch by sampling from a Poisson distribution with mean equals *c · t*, where *t* is the scaled branch length and *c* is the event multiplier that controls the number of CNAs at the leaves of the tree. To mimic the punctuated mode of tumor evolution as in [10], at the edge to the root, we sampled a multiplier *m* from a Poisson distribution with mean *λ* = 8 and sampled the number of CNA from a Poisson distribution with mean *m · c · t*. To evaluate how the complexity of copy number events can affect the performance of each method, we set *c* to 90 and 250, corresponding to the cases of low and high frequency of copy number events, respectively. The size of each CNA event is determined by sampling from an exponential distribution with mean=10Mbp, plus a minimum CNA size of 2Mbp. Whether the CNA resulted in a gain or a loss of copy is determined by sampling from a binomial distribution with *p* = 0.5. Then, to determine the location of each CNA event, we first randomly sample the allele on which the CNA is going to occur from the paternal and maternal alleles with a binomial distribution with *p* = 0.5. Then, we linearize the genome and sample the genomic coordinate *x* where the CNA occurs from a probability distribution whose density function is given by 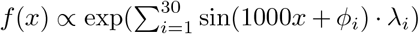. We fixed *ϕ*_*i*_ *∼ Uniform*(−*π, π*) and *λ*_*i*_ *∼ Uniform*(0, *a*), where *a* is a user-specified parameter that controls the non-uniformity of the distribution. Using a normalized sum of sines with a random phase as the distribution to sample copy number events provides enough randomness and allows control of overlap during simulation [20]. For the purpose of this study, we set *a* = 0.6 as we found the CNAs simulated under such a setting best resembled what we observed from the biological data sets. On average, more than 90% of the CNAs overlapped with at least one other CNA. After all CNA events are added along the branches, the genome is divided into 15k non-overlapping bins, and the copy number of each bin in a leaf is calculated from all copy number events along the path from the root to the leaf.

### 2.3 Running the Clustering Algorithms

We ran each clustering algorithm on dimensions that were reduced through Principal Component Analysis (PCA) to reduce the data to a 50-dimensional space and t-distributed Stochastic Neighbor Embedding (t-SNE) to a 2-dimensional space, in addition to the original dimensions. We then ran each clustering algorithm across a specified range of parameters (see details below) to determine the optimal parameter. The silhouette score [26], a metric used to assess the quality of cluster assignments by measuring how similar an object is to its own cluster compared to other clusters, was computed for the generated cluster assignment. We then selected the parameter that resulted in the highest silhouette score for the respective combination of the clustering method and dimensionality-reduction technique applied to the data set, with the only exception being when we evaluated the performance with respect to the dimension-reduction technique. We ran the Louvain algorithm by inputting the K nearest neighbors (KNN) graph constructed from the copy number profile matrix. Specifically, we first calculated a pairwise Euclidean distance matrix on the original dimensional space for all cells with [25] and then constructed the input network by connecting each cell to its K nearest neighbors. We varied K from 5 to 100. We ran K-means with the implementation in [25]. We varied K, the number of clusters from 4 to 50 given our observation from the biological data set. We ran hierarchical clustering with the implementation in [25] with the Euclidean distance as metric and *Avergae* as linkage. We varied K, the number of clusters, from 4 to 50.

### 2.4 Clonal Discordance Score

In order to evaluate the consistency between clustering results and the phylogenetic tree structure, we use the clonal discordance score as suggested in [29]. To calculate this score, we first assign the cluster label of each leaf as the profile of that leaf in the tree. Next, we use the implementation in [33] to compute the parsimony score of the tree. Here, all label transitions are treated as substitutions with a uniform weight of one. The minimum number of label transitions that can be expected for a set of *n* distinct cluster labels within a tree is *n*− 1. The clonal discordance score is then calculated by subtracting the minimum expected number of label transitions (*n* − 1) from the actual number of label transitions. This adjustment helps account for the effect of the number of clusters, thus providing an unbiased metric of how well the cluster assignments match with the phylogenetic tree.

### 2.5 Normalized Entropy of Clustering

Balanced cluster assignment ideally has nearly equal data points in every cluster, while imbalances may suggest overlooked data patterns. To evaluate the balances of cluster assignments, we computed the normalized entropy [30]. Specifically, for a clustering assignment with *k* clusters, the probability of a single cell belonging to a specific cluster, *P* (*i*), can be calculated by the ratio of cells in cluster *i* to the total cells. The entropy for cluster then *i* is *H*(*i*) = −*P* (*i*) *·* log_2_ *P* (*i*)and the total entropy is 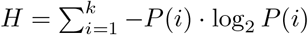.

For standardization, the entropy is normalized against the maximum possible entropy, which occurs when clusters are of equal size. Specifically, the maximum entropy for *k* clusters can be computed by 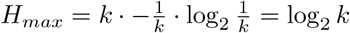. The normalized entropy is thus 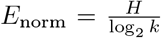, with values closer to 1 indicating a more balanced distribution across clusters.

### 2.6 Triplet Distance between Trees

We assess the accuracy of the inferred clonal trees and full binary tree compared to the ground truth trees in simulation by employing triplet distance [7], using the implementation in [28]. The triplet distance evaluates the concordance of tree topologies based on the arrangement of leaf nodes within triplet structures, applicable when comparing both clonal trees with full binary trees and between two full binary trees. For every set of three leaves in a rooted tree, termed a *triplet*, there exist three possible binary tree topologies. To calculate the triplet distance between two phylogenetic trees, we counted the number of different phylogenetic relationships among all possible sets of triplets in the trees. Specifically, this is accomplished by determining whether the binary tree topology induced by those triplets in the two phylogenetic trees is identical or not. Note that for clonal trees, an unresolved triplet (in the case that three leaves belong to the same cluster) is also counted as a difference, as it does not match the fully resolved counterpart in a full binary tree. We then normalize the triplet distance by dividing the number of disagreeing (unresolved) triplets by the total number of triplets in the tree. This metric reflects the number of triplet structures that differ or are unresolved compared to a binary tree. It provides a clear measure of their topological divergence.

## 3 Results

On the simulated data, we evaluated each of the approach’s steps individually in order to isolate their strengths and weaknesses. We evaluated the impact of the chosen dimensionality reduction and clustering algorithms on runtime and accuracy. Rather than assuming a true clustering for benchmarking, we evaluated the quality of the clusters in terms of their agreement with the tree on which the cells evolved. We also assessed the accuracy of the AP-derived CRPs against traditional consensus profile (CP) derived CRPs, in terms of the goodness of their representation of the CNA profiles of all cells within that cluster. Finally, we studied the accuracy of the final tree obtained by NestedBD.

For the biological data, since the ground truth is unknown, we investigated the disagreements in the results obtained by different methods.

### 3.1 Performance on Simulated Datasets

In the first experiment, we assessed the performance of three clustering algorithms—Louvain, K-means, and hierarchical clustering—on clustering single cells based on their CNA profiles, using simulated datasets with 1000, 5000, and 10000 cells. The datasets were simulated with “high” and “low” numbers of CNA events. The average normalized pairwise Manhattan distances between simulated single cells in data simulated under these two settings are 0.76 and 0.39, respectively. More details of the simulation setup are available in Section ‘Simulation Protocol.’ To examine the effects of dimensionality reduction on clustering accuracy and runtime efficiency, we employed PCA with 50 dimensions and t-SNE with two dimensions. The choice of number of dimensions to be used for each technique is selected so as to follow the settings in empirical studies [24,34,16]. We also performed clustering on the original, high-dimensional data. This approach allowed us to directly compare and understand the impact of dimensionality reduction on the clustering algorithms. The specific configurations used for each method can be found in Section ‘Running the Clustering Algorithms.’

#### Clustering Runtime

Table 1 summarizes the runtime for each combination of clustering algorithm and dimensionality reduction technique. Applying PCA significantly reduced the runtime for all clustering methods across all data sizes. While t-SNE also reduced the runtime of clustering compared to scenarios without dimensionality reduction, its performance was not as effective as PCA’s. The runtime for clustering after t-SNE dimensionality reduction was notably higher than that after PCA, particularly in larger datasets. This increase in runtime with t-SNE can be attributed to the fact that although computing distances in the lower-dimensional space is generally faster, t-SNE itself scales poorly with the number of samples. As a result, while t-SNE may have certain advantages in preserving local structure [13], PCA is a particularly effective technique for reducing runtime.

**Table 1.** Runtime of clustering algorithms with dimensionality reduction. Runtime for Louvain, K-means, and Hierarchical Clustering methods on single-cell CNA data of different sizes, both with and without dimensionality reduction techniques. Dimensionality reduction is implemented using PCA with 50 dimensions and t-SNE with two dimensions. The average and standard deviation of runtime for each scenario are summarized on 10 replicates.

| Number of Cells | Dimensionality Reduction | Total Time Taken for Dimensionality Reduction and Clustering (seconds) |  |  |
| --- | --- | --- | --- | --- |
|  |  | Louvain | K-means | Hierarchical |
| 1000 | None | 49.73 ± 8.20 | 46.15 ± 1.35 | 15.54 ± 2.36 |
| 1000 | t-SNE | 26.67 ± 4.06 | 26.58 ± 4.05 | 26.46 ± 4.06 |
| 1000 | PCA | 1.00 ± 0.09 | 0.99 ± 0.08 | 0.68 ± 0.08 |
| 5000 | None | 815.29 ± 133.90 | 249.70 ± 21.33 | 299.61 ± 53.88 |
| 5000 | t-SNE | 414.82 ± 69.85 | 411.61 ± 69.86 | 412.07 ± 69.83 |
| 5000 | PCA | 8.00 ± 0.48 | 3.29 ± 0.16 | 3.39 ± 0.14 |
| 10,000 | None | 3197.50 ± 494.54 | 517.77 ± 9.38 | 1238.30 ± 187.19 |
| 10,000 | t-SNE | 1593.66 ± 228.15 | 1581.03 ± 228.29 | 1584.12 ± 228.10 |
| 10,000 | PCA | 25.50 ± 2.32 | 4.56 ± 0.17 | 9.12 ± 0.51 |

In terms of comparing the clustering methods, we observed that without dimensionality reduction, hierarchical clustering was the fastest method for the smallest dataset size with 1,000 cells. However, its relative efficiency decreased with increasing dataset size, becoming the slowest among the three methods for the largest datasets (10,000 cells). K-means, on the other hand, consistently showed moderate runtimes across all scenarios, striking a balance between speed and scalability. Louvain, being the slowest for the smallest dataset without dimensionality reduction, exhibited significant runtime improvements with dimensionality reduction, particularly with PCA, suggesting its scalability benefits from dimensionality reduction more markedly than the other methods. The increase in runtime with data size is evident for all methods and scenarios, but the impact of dimensionality reduction in mitigating this increase is significant. PCA allows for a more scalable approach to clustering large datasets, as seen in its dramatic reduction of runtimes across all clustering methods and data sizes.

#### Clustering Accuracy

While the ground truth is known in simulations, what constitutes a true cluster is not easy to define [12]. Therefore, in assessing the accuracy of clustering methods here, rather than assuming knowledge of true clusters, we quantify the consistency of the obtained clusters with the trees on which the genomic profiles evolved. For this purpose, we used the clonal discordance score [29]. This score reflects the minimum number of cluster label transitions needed to align with the phylogeny’s cell arrangement, where a higher score signifies lower concordance and more transitions required (more details in Section ‘Clonal Discordance Score’). The score serves as an effective metric to assess and compare the reliability of clustering and dimensionality reduction methods in the initial stage of phylogenetic tree inference without the need to designate a ground truth clustering. Results are summarized in Fig. 2 and table 2.

**Table 2.** Comparison of Clustering Methods Based on Clonal Discordance Scores. The table summarizes the t-test results of the clonal discordance scores for cluster assignments generated by Louvain, K-means, and Hierarchical Clustering across 10 simulated replicates. Scores are ranked by the median of 10 replicates, and p-values are computed to compare the best method against the second best, assessing significant differences. The labels “high” and “low” categorize the extent of CNAs in the simulated data (see Section Simulation Protocol).

| Number of Cells | Extent of CNA events | Best Method | p value from t-TEST |
| --- | --- | --- | --- |
| 1kleaf | High | Louvain | 0.00235 |
|  | Low | Louvain | 0.04099 |
| 5kleaf | High | K-Means | 0.01112 |
|  | Low | K-Means | 0.01781 |
| 10kleaf | High | K-Means | 0.01276 |
|  | Low | K-Means | 0.00921 |

**Fig. 2.**
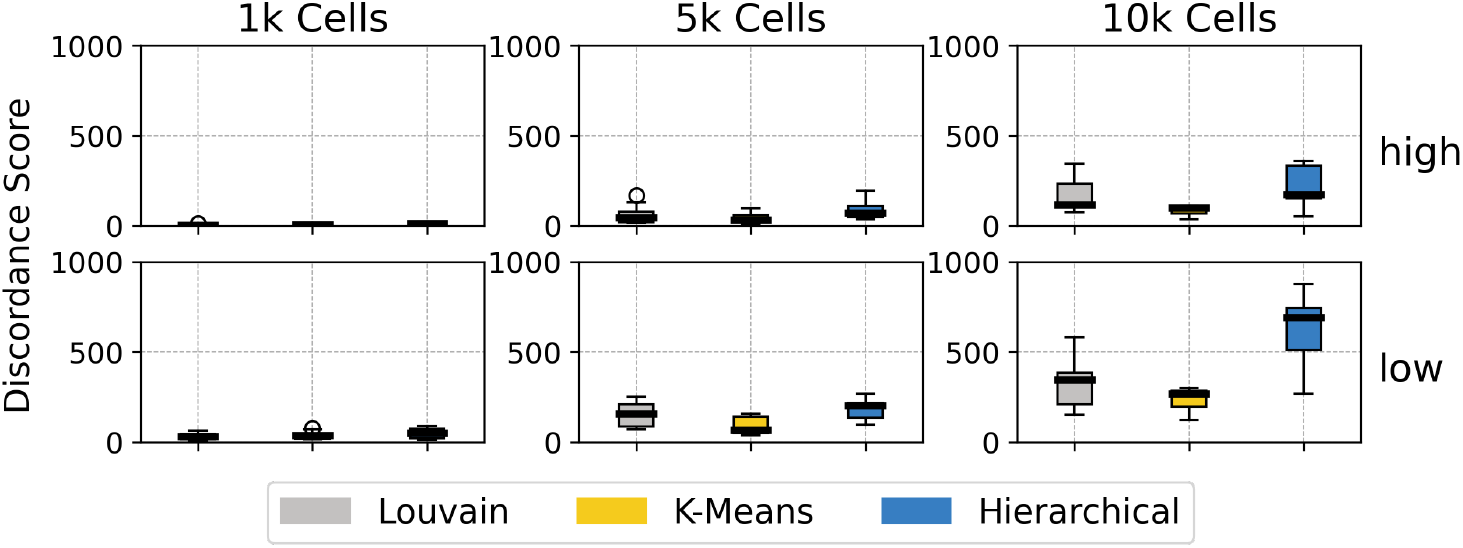
Clustering Performance on Simulated Data: Clonal Discordance. The box plots show the distributions of the clonal discordance scores for cluster assignments generated by Louvain, K-means, and Hierarchical Clustering. The results are summarized on 10 simulated replicates. “high” and “low” refer to the extent of CNAs in the simulated data (see Section ‘Simulation Protocol’).

The results show that Louvain performs better with datasets with fewer cells; however, its performance declines significantly with increased cell count. K-means, on the other hand, shows relatively consistent accuracy across all data sizes, even with large datasets of up to 10k cells, establishing itself as a robust choice. The accuracy of hierarchical clustering was consistently lower compared to the other two methods, suggesting that it may not be the preferred method in this particular context. We also observed that the accuracy of clusters is poorer on data with a lower amount of CNA events. This is potentially due to insufficient signal in the datasets with low CNA event counts.

We also studied the number and balance of generated clusters (Fig. 3 and Fig. 4). We observed that the Louvain clustering method consistently resulted in a higher number of clusters than K-means and hierarchical clustering. Moreover, the number of clusters generated by Louvain clustering increased with the increase in the number of cells. On the other hand, K-means and hierarchical clustering consistently produced lower numbers of clusters, regardless of the number of cells.

**Fig. 3.**
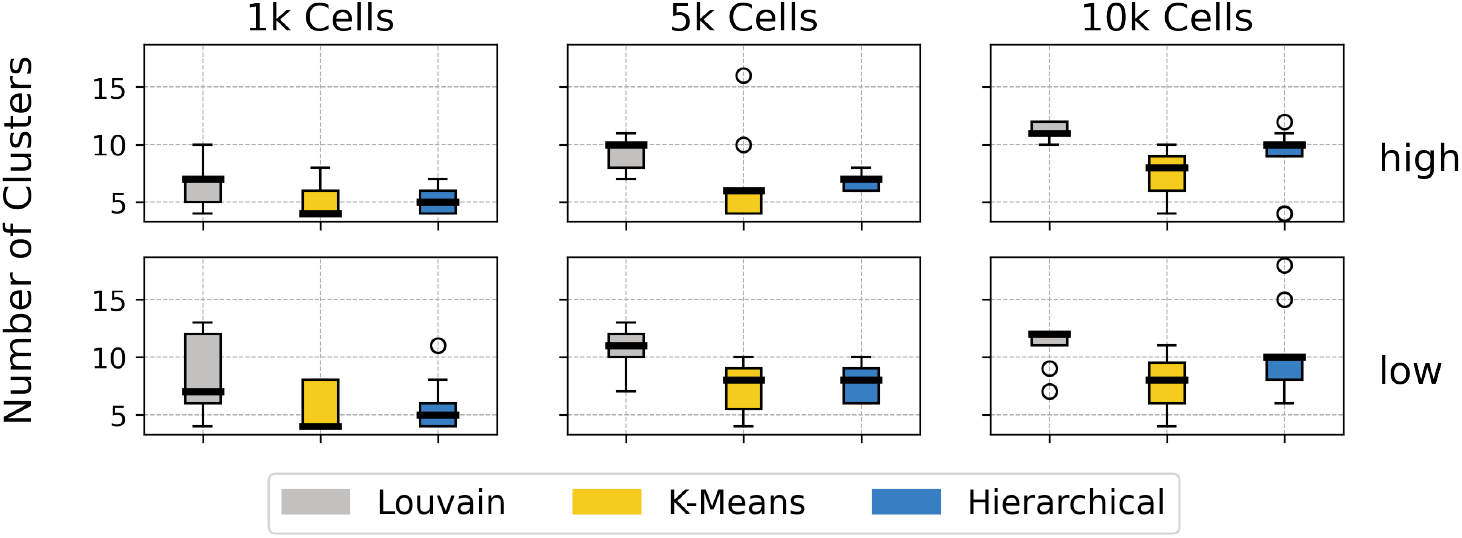
Clustering Performance on Simulated Data: Number of Clusters. The box plots show the distributions of the number of clusters generated by Louvain, K-means, and Hierarchical Clustering on simulated data. The results are summarized on 10 simulated replicates. “high” and “low” refer to the extent of CNAs in the simulated data (see Section ‘Simulation Protocol’).

**Fig. 4.**
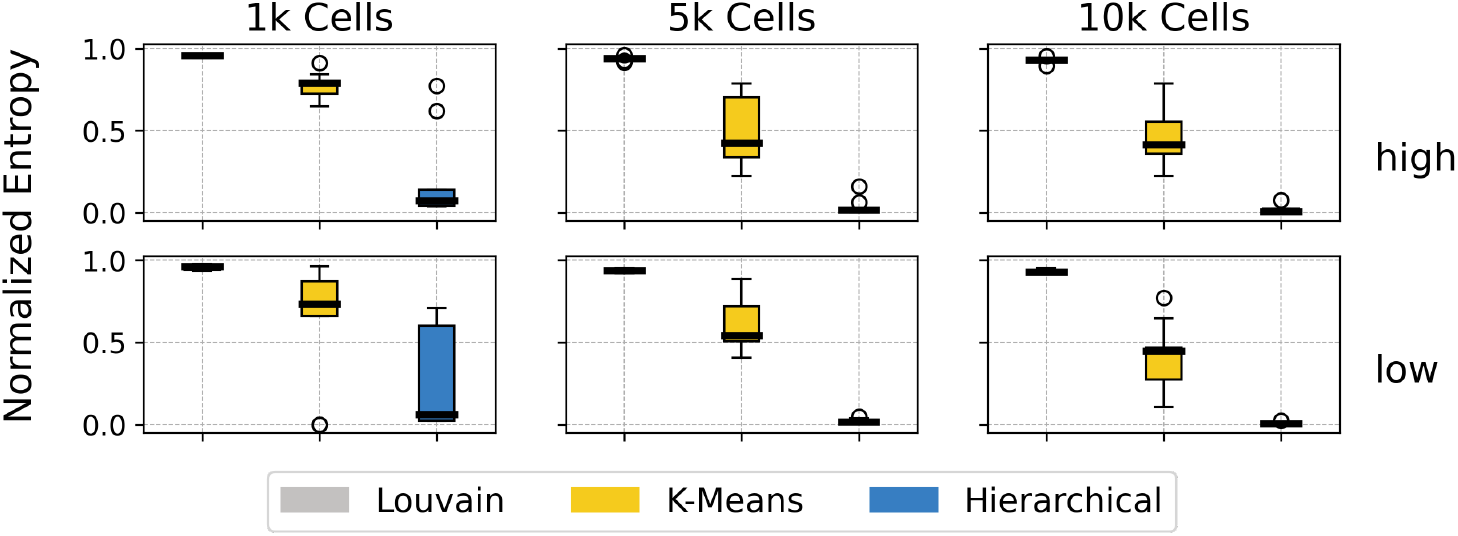
Clustering Performance on Simulated Data: Cluster Balance. The box plots show the distributions of the normalized entropy of cluster assignments generated by Louvain, K-means, and Hierarchical Clustering. The results are summarized on 10 simulated replicates. “high” and “low” refer to the extent of CNAs in the simulated data (see Section ‘Simulation Protocol’).

When evaluating the balance of different clustering methods using normalized entropy, a measure that quantifies the uniformity of cluster size distribution across a dataset (see details in Section ‘Clonal Discordance Score’), we found that Louvain clustering generated a uniform cluster distribution. In comparison, K-means showed variability in cluster sizes. Hierarchical clustering produced a notably different cluster size distribution, characterized by significantly unbalanced clusters, including many small or even singleton clusters, with the normalized entropy value approaching zero. These results, coupled with the ones in Fig. 2, indicate that clusters generated by Louvain are best, whereas those generated by hierarchical clustering align the least with the true tree.

We summarize the impact of using dimensionality reduction before running the clustering algorithm on the accuracy of cluster assignments in Fig. 5. While we observed better runtime efficiency with the use of dimensionality reduction techniques under some scenarios, as detailed in Table 1, our analysis also revealed decreased clustering accuracy. Specifically, clustering performed on data in its original dimensional space yielded results more closely aligned with the true evolutionary trees, implying a trade-off between efficiency and the retention of critical signals for accurate cluster identification.

**Fig. 5.**
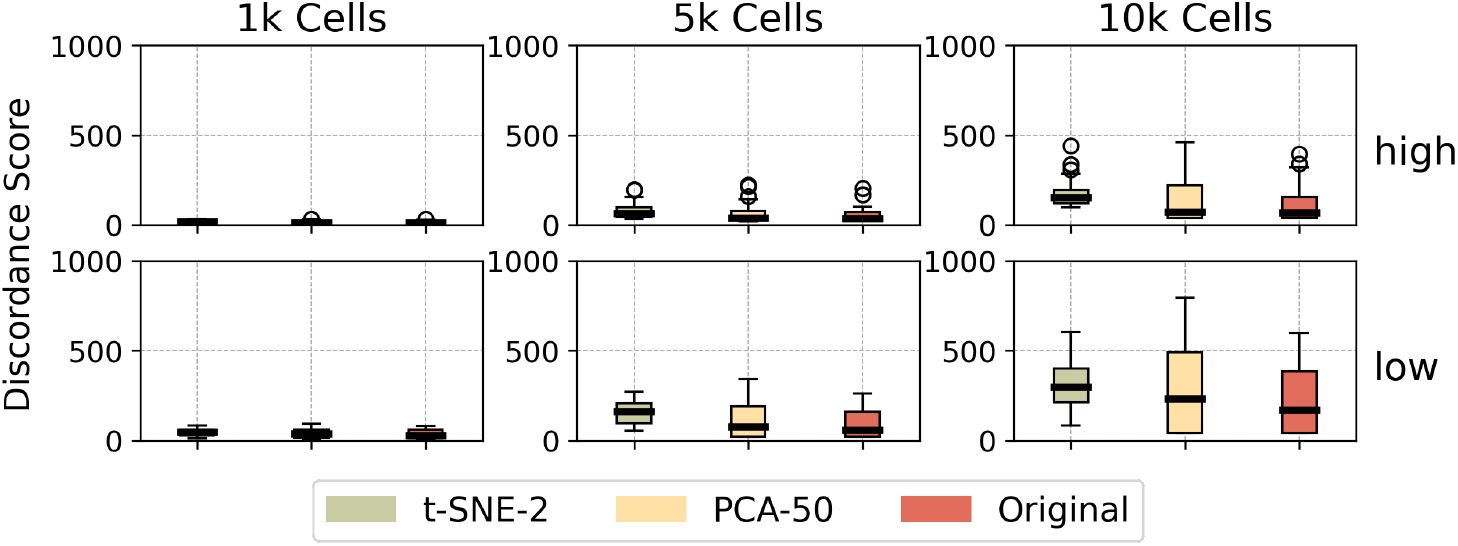
Clustering Performance on Simulated Data: The impact of dimensionality reduction. The box plots show the distributions of clonal discordance scores for cluster assignments using the original dimension and reduced dimensions (via t-SNE and PCA). The results are summarized for all clustering methods on 10 simulated replicates. “high” and “low” refer to the extent of CNAs in the simulated data sets.

#### Quality of Cluster-Representative Profiles (CRPs)

After computing the cluster assignments of cells, we generated a CRP for each cluster. Unlike existing studies that compute consensus profiles, we propose ancestral profile (AP) inference on an evolutionary tree of all copy number profiles in the cluster and using the profile at the root of the tree as the CRP for that cluster. In this work, we assessed the performance of this AP-based approach using neighbor-joining (NJ) [27] for constructing an evolutionary tree from all cell profiles within a cluster and maximum parsimony (MP) for AP inference. In order to obtain a rooted NJ tree, we introduced a diploid copy number profile to every cluster and treated it as the outgroup at which we rooted the tree. In addition to the NJ tree, we also inferred the ancestral profile on the true tree, which is known in simulated data, in order to quantify the effect of tree estimation error on obtaining the CRP via ancestral reconstruction. We benchmarked the performance of the AP-based approach against the traditional CP-based approaches, which derive the CRP from the mean, median, and mode of copy numbers across all cells in the cluster.

In cases where a cluster does not align perfectly with the ground truth tree (i.e., the cluster does not constitute a clade in the tree), we reconstructed an ancestral profile for each cluster using the sub-tree containing all cells in the cluster. To determine the accuracy of the inferred CRPs, we calculated the Hamming distance and L1 norm between the inferred CRP and the true profile of the lowest common ancestor (LCA) of all copy number profiles in a cluster. The results are summarized in Fig. 6.

**Fig. 6.**
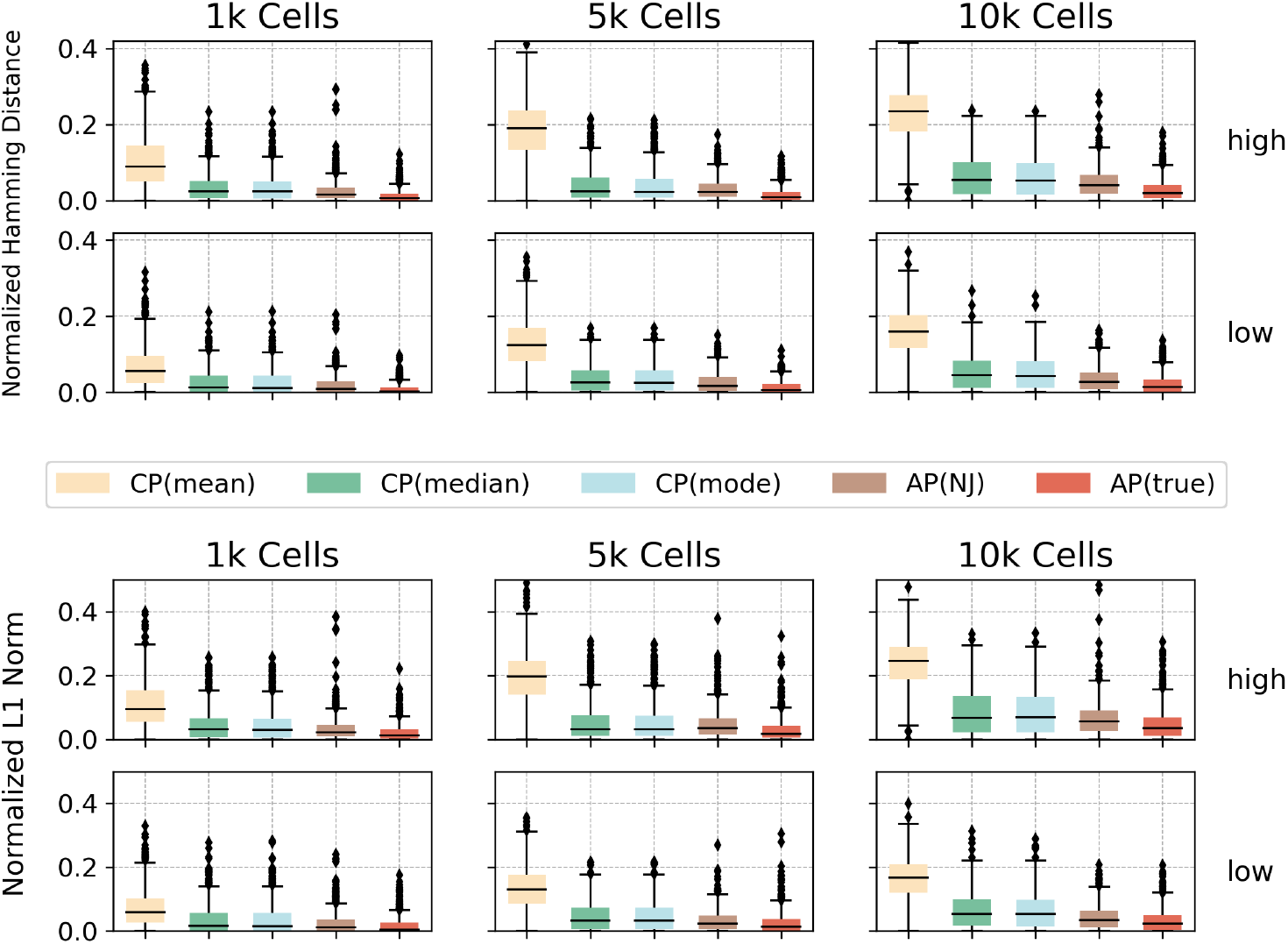
Accuracy of CRPs. The box plots show the distributions of CRP estimation error measured by the normalized Hamming distance and L1 norm between the true and inferred CRP of a cluster. The inference methods used include three consensus profile (CP) methods — mean, median, mode — and two ancestral profile (AP) methods using the NJ and true tree. “high” and “low” correspond to the extent of CNAs in the simulated data (see Section ‘Simulation Protocol’).

The results show that among the three consensus profiles, the median and mode are more reliable than the mean. However, we found that ancestral profiles inferred from the NJ tree were more accurate than these consensus profiles. The accuracy of profile inference decreases as the number of cells in a cluster increases, regardless of the profile inference method used. This observation aligns with the one about decreased accuracy of clusters with larger numbers of cells we discussed above. The extent of CNA events has little effect on accuracy. Even though cluster quality is lower for low counts of CNA events (Fig. 2), the quality of the inference CRPs is better than those inferred under the higher extent of CNAs. This could be due to the small number of events, which gives rise to a simpler CRP inference, even if the cluster itself is not very accurate.

#### Quality of Evolutionary Trees Inferred on CRPs

An important goal of large-scale evolutionary analysis of scDNA data is to obtain an evolutionary, or mutation, tree on the clusters (Step (6) in Fig. 1). Here we propose to infer a tree on the CRPs using the NestedBD method [20]. It is important to note that NestedBD is a fully Bayesian method and is not scalable to thousands or tens of thousands of cells. Furthermore, we inferred an NJ tree, using both pairwise Hamming distances and pairwise ZCNT distances as in [29], on the full dataset to assess the quality of evolutionary inference without adopting the clustering pipeline. We also attempted to run MEDICC2 [14] on the full data; however, its computation was not completed within a reasonable time frame for this experiment and was therefore excluded.

To evaluate the accuracy of the inferred trees, we calculated the triplet distance [7], which serves as a robust measurement of topological agreement between the cluster tree or full binary tree and the ground truth tree (see Section ‘Triplet Distance between Trees’). The runtimes of tree inference methods are summarized in Table 3. Two observations are in order. First, NJ is fast enough to be run on large datasets without clustering. Second, clustering allows a fully Bayesian method like NestedBD to be applicable on such large datasets.

**Table 3.** Runtime of Tree Inference Methods. NJ trees were inferred on the full datasets using the ZCNT and Hamming distances, whereas the results in this table for NestedBD are based on using the CRPs obtained from consensus profiles. The mean and standard deviation of runtimes for each simulation setting are summarized on 10 replicates.

| Number of Cells | Total Time Taken for each Inference Method (in seconds) |  |  |  |  |
| --- | --- | --- | --- | --- | --- |
|  | NJ (ZCNT) | NJ (Hamming) | NestedBD (Louvain) | NestedBD (K-means) | NestedBD (Hierarchical) |
| 1,000 | 192.80 | 19.63 | 88.30 | 81.08 | 92.90 |
| | $\pm 7.06$ | $\pm 2.88$ | $\pm 10.31$ | $\pm 8.73$ | $\pm 3.97$ |
| 5,000 | 6025.84 | 1628.20 | 956.46 | 297.06 | 397.06 |
| | $\pm 1300.22$ | $\pm 275.77$ | $\pm 110.67$ | $\pm 52.41$ | $\pm 60.00$ |
| 10,000 | 35685.76 | 13580.17 | 3345.74 | 526.03 | 1185.13 |
| | $\pm 5726.97$ | $\pm 1807.60$ | $\pm 482.33$ | $\pm 100.98$ | $\pm 171.13$ |

The accuracy of the inferred trees is summarized in Fig. 7 and Fig. 8. As the results show, NestedBD trees inferred on CRPs computed via ancestral profiles are consistently the most accurate among all methods. While NJ is able to run on large datasets, our results show that clustering and inferring trees on clusters would provide more accurate trees. Fig. 8 shows that the accuracy of the clusters has a big impact on the quality of the trees inferred from the CRPs. In summary, these results demonstrate that the pipeline outlined in Fig. 1 is effective for inferring evolutionary trees on large single-cell data especially when clustering results are accurate and CRPs using ancestral profiles are used.

**Fig. 7.**
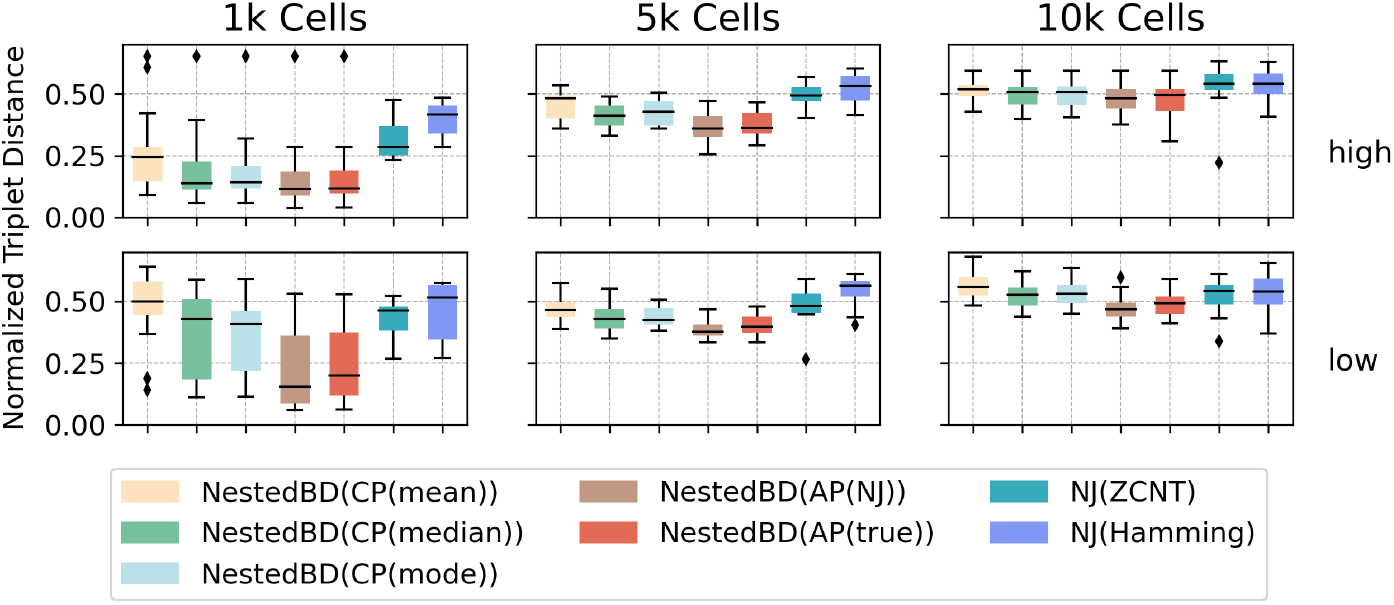
Inferred Tree Accuracy. The box plots summarize the distributions of normalized triplet distances between the ground truth tree and the inferred trees. Trees were inferred by NJ on the full datasets using two measures of pairwise distances— ZCNT and Hamming—and by NestedBD on the CRPs of Louvain, K-means, and hierarchical clustering clusters computed in five different ways (see main text). “high” and “low” correspond to the extent of CNAs in the simulated data (see Section ‘Simulation Protocol’).

**Fig. 8.**
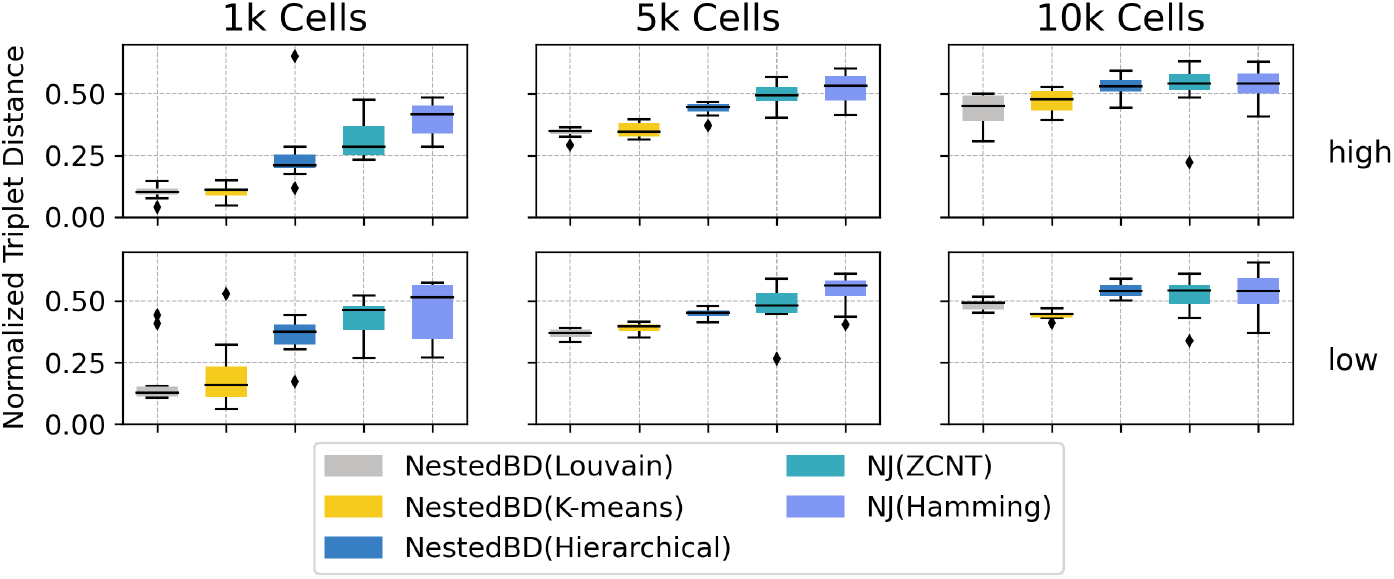
Inferred Tree Accuracy by Clustering Method. The box plots summarize the distributions of normalized triplet distances comparing the ground truth tree with inferred trees. Trees were inferred by NJ on the full datasets using two measures of pairwise distances—ZCNT and Hamming—and by NestedBD on the CRPs using AP(NJ) on the clusters obtained by all three clustering methods. “high” and “low” correspond to the extent of CNAs in the simulated data (see Section ‘Simulation Protocol’).

### 3.2 Performance on Biological Data

We obtained single-cell whole-genome sequencing (WGS) data from [9] and selected the copy number profile data from three different samples, each with varying numbers of single cells. Specifically, these samples include SA1050 with CNA profiles from 990 cells, SA501 with CNA profiles from 2473 cells, and SA906b with CNA profiles of 5716 cells. Similar to the simulated data, we applied the three clustering algorithms, Louvain, K-means, and hierarchical clustering, to the CNA profile data from each sample, without applying dimensionality reduction as our results indicated they negatively impact the clustering quality. A visualization of the clustering results on the data is given in Supplemental Fig. S 1. The specific configuration used for each method can be found in Section ‘Running the Clustering Algorithms.’

Since the ground truth is unknown for empirical datasets, we evaluated the different methods using two approaches. First, we assessed the agreement among the clusters generated by each method by computing the Adjusted Rand Index (ARI), where higher ARI values indicate greater agreements among the clustering. Second, we evaluated the agreement of evolutionary trees constructed from CRPs by computing the triplet distance between them, with a lower triplet distance suggesting a closer agreement between the trees.

#### Disagreement Among Clustering Methods

We plotted the pairwise ARI values for pairs of methods on the three samples, summarized at the left of Fig. 9. The results show that Louvain and K-means have the highest agreement, whereas hierarchical clustering produces clusters that are most different from the other two methods. Furthermore, the agreements between methods decrease with the increase in the size of the data in each sample (SA1050 has the smallest number of cells and SA906b has the largest). These results match our observations on simulated data where Louvain and K-means have the highest accuracy and the accuracy of all methods dropped with increasing data sizes. These results further confirm our findings on simulated data that clustering methods could give very different results and simply marking an arbitrary choice of the method might not be a safe practice.

**Fig. 9.**
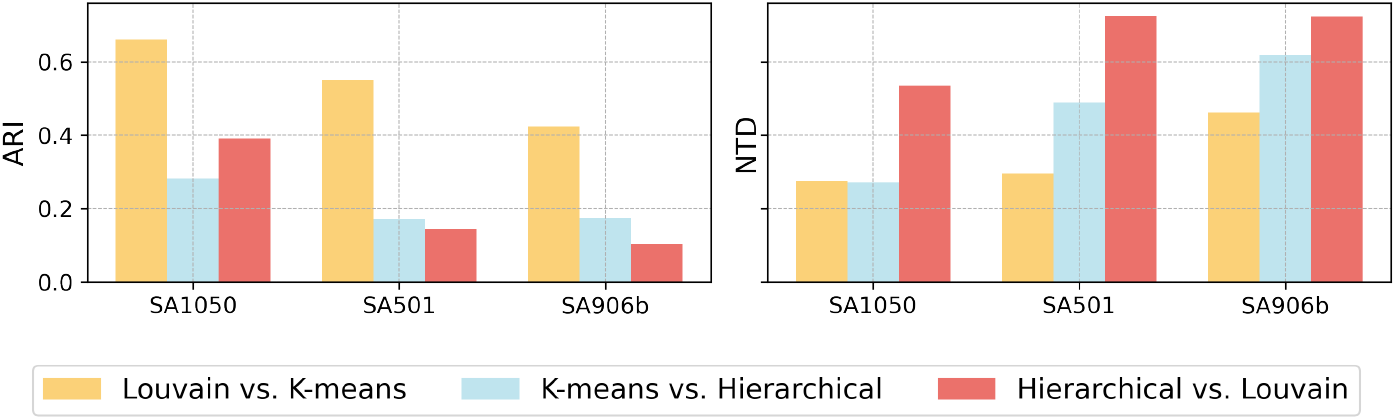
Adjusted Rand Index (ARI) values and Normalized Triplet Distances (NTD). (Left) ARI’s were calculated on the clusters obtained by pairs of methods on the data from three samples. (Right) NTD between pairs of trees inferred on the CRPs of clustered data using three clustering methods in each sample. Each CRP is the ancestral profile of the NJ tree on a cluster.

#### Disagreement Among Inferred Trees

Following the clustering process, we used NestedBD [20] to infer evolutionary trees for each sample dataset, utilizing CRPs based on the ancestral profiles computed on NJ trees. The pairwise normalized triplet distances between trees inferred on clusters produced by the three clustering methods are summarized in the right of Fig. 9.

Our results indicate a clear trend linking the quality of clustering to the accuracy of the resulting trees. Specifically, lower ARI values between clusters correlated with higher normalized triplet distances between trees, particularly as the number of cells increases, indicating that the quality of trees is directly associated with the similarity of the clusters from which they were derived. This correlation is further supported by the consistent clustering produced by Louvain and K-means algorithms (Fig. 9 left), which also led to the generation of closely matching phylogenetic trees.

Comparing hierarchical clustering with the other two methods shows a similar trend, where a drop in clustering accuracy is linked to an increase in tree distance. However, this correlation became less clear on the largest sample, suggesting that the impact on tree inference is more attributable to constraints imposed by the tree inference algorithms rather than the variances in clustering itself, especially in cases of significant clustering disagreement. This pattern further emphasizes the importance of using accurate tree inference methods.

## 4 Discussion

Evolutionary analysis of large scDNAseq datasets (thousands or tens of thousands of individual cells) is limited by the scalability of the employed methods. In this paper, we introduced a divide-and-conquer approach that is flexible enough to scale up existing and newly developed scDNAseq evolutionary analysis methods. We assessed the performance of the approach on both simulated and biological data while experimenting with a variety of methods for the various steps of the approach. As we mentioned above, the goal of our study was not to identify the best clustering, dimensionality reduction, or tree inference method; rather, it was to demonstrate the flexibility and effectiveness of the approach.

Standard phylogenetic methods, like NJ, are scalable to thousands of single cells. However, we found their accuracy to be lowest as compared to ones obtained by our approach and using CNA-specific methods, such as NestedBD. It is worth mentioning that NestedBD is not scalable to the datasets we experimented on in this study.

An important finding from our study is the large discrepancy among results based on different clustering techniques. In a typical phylogenetic supertree implementation [36], the “divide” step is not based on clustering methods. A much more accurate method in that domain is to first estimate a tree on the full dataset using a scalable, but not necessarily very accurate, method, and use the tree to guide the division of the dataset. Furthermore, while we experimented with data partitioning via clustering, in phylogenetic supertree methods, the subsets are overlapping in order to glue the subtrees properly. Studying the performance of both techniques (a guide tree and overlapping subsets) in our general approach is a direction for future research.

## Acknowledgments

This research was supported in part by the National Science Foundation, grants IIS-1812822 and IIS-2106837 (L.N.).

